# Quantifying colour difference in animals with variable patterning

**DOI:** 10.1101/859074

**Authors:** Tereza Dračková, Radovan Smolinský, Zuzana Hiadlovská, Matej Dolinay, Natália Martínková

**Affiliations:** Institute of Biostatistics and Analyses, Masaryk University, Kamenice 3, 62500 Brno, Czech Republic; Department of Biology, Faculty of Education, Masaryk University, Poříčí 7, 60300 Brno, Czech Republic; Institute of Animal Physiology and Genetics, Czech Academy of Sciences, Veveří 97, 60200 Brno, Czech Republic; Institute of Vertebrate Biology, Czech Academy of Sciences, Květná 8, 60365 Brno, Czech Republic; Department of Botany and Zoology, Masaryk University, Kotlářska 2, 61137 Brno, Czech Republic; RECETOX, Masaryk University, Kotlářska 2, 61137 Brno, Czech Republic

**Keywords:** colouration, Reptilia, image analysis, colour pattern, RGB, CIELAB

## Abstract

Colour pattern influences behaviour and modifies survival of organisms through perception of light reflectance. Spectrophotometric methods used to study colour navigate between precision and accuracy of reflectance across wavelengths, while photographic methods are generally used to assess the complexity of colour patterns. Here, we compare how colours characterised using point measurements (simulating spectrophotometry) on the skin of a sand lizard (*Lacerta agilis*) differ from colours estimated by clustering pixels in a photograph of the lizard’s body. We found that point measurements adequately represented the dominant colour of the lizard; however, where colour patterning influenced measurement geometry, image analysis outperformed point measurement as regards stability between technical replicates on the same animal. The greater colour variation established from point measurements increased further under controlled laboratory illumination. Both methods revealed lateral colour asymmetry in sand lizards. We conclude that studies assessing the impact of colour on animal ecology and behaviour should utilise hyperspectral imaging, followed by image analysis that encompasses the whole colour pattern.

## Introduction

Studies using spectrophotometric analysis have shown that organisms differ in their ability to recognise and perceive light of different wavelengths, specifically through the triggering of signal protein expression in the photoreceptor cells (Testylier and Gourmelon 1987). To calibrate the spectrophotometer, measurements of a standard and sample need to be set to within a millimetre of their relative positions, and to within a degree of the angle between the light source and the detector (Johnsen 2016). The precise geometry of the measurement poses significant technical difficulties as standards, unlike live animals, are usually flat, while animals tend to resist immobilisation, not least by breathing. In relation to these technical constraints, spectrophotometric analysis during ecological research is most successful when using skin derivatives that can be removed from the animal (e.g. feathers, scales) or flat appendages (e.g. insect wings, fish fins) (McCoy and Prum 2019). In comparison, the challenges of spectrophotometry on live animals or on objects with informative colour patterns are prohibitive and, as such, colour pattern complexity has been understudied (but see Maia et al. 2019; Šulc et al. 2019).

In this study, we set out to examine colour variation in sand lizards (*Lacerta agilis*, Reptilia: Lacertidae), a species for which use of spectrophotometry has proved problematic. Sand lizards are relatively small (total length up to 240 mm), exhibit sexual dimorphism and sympatric colour forms result in variation across the Palearctic (Bischoff 1988; Andres et al. 2014). These colour forms differ both in predominant colour and in the position and number of stripes and spots on the animal’s body (Kotenko and Sviri-denko 2010). In addition to colour variation and sexual dimorphism, sand lizard colour changes with the seasons. For example, spots contributing to the colour pattern enlarge during the reproduction season (spring and early summer), and males are brightly coloured in the spring and become duller as the seasons progress. Depending on the colour form, males may display hues of green and/or brown during the mating season, while females and juveniles display hues of brown throughout the year. In some regions, e.g. Iberia, the scales of sand lizards reflect UV light, while in other regions, e.g. Scandinavia, sand lizards do not possess this characteristic (Font and Pérezi de Lanuza 2007).

Owing to the difficulties of undertaking spectrophotometry on small, living animals with complex colour patterns, we decided to quantify the influence of model characteristics by point measurement. In doing so, we compare two alternative methods for quantifying colour differences from standardised photographs. The first simulates the point measurements of a spectrophotometer, with measurements taken at body regions defined by the animal’s morphology, regardless of any specific colour pattern; while the second assesses a complete photograph and sorts individual pixels by colour, irrespective of colour pattern. In addition, we compare two alternative lighting conditions, i.e. fluorescent darkroom bulbs vs. natural environmental light, in order to evaluate the stability of the colours estimated. Our overall aim was to estimate the degree to which colour pattern influences accuracy of colour measurement.

## Materials and methods

### Sampling

We sampled two different sand lizard populations during the sand lizard’s active season, the first from an orchard in Unín (48.72 N, 17.24 E) in Slovakia between April and September 2007 and the second from an orchard in Hustopeče (48.93 N, 16.72 E) in the Czech Republic between May and August 2018. The disparity in the length of sampling season was caused by differing weather conditions in 2007 and 2018. Both localities were screened before and after the collection periods, allowing us to reliably confirm the first and last days when active animals were observed. All lizards were caught by hand or by noosing at weekly intervals and each was individually marked by toe clipping (Waichman 1992).

To estimate the age of individual animals, we used tpsDig v.2.02 (Rohlf 2005) software to measure body length from rostrum to anus (*L*) from digital images. Lizards over 6 cm were considered adults, specimens of 4.7 to 6 cm were considered sub-adults that had over-wintered once but had not yet reproduced and those smaller than 4.7 cm were considered juveniles that had hatched in the given season (adjusted after Bischoff 1984). Lizards caught in 2007 were transported to the laboratory, where they were photographed in a darkroom using standard light conditions and released the next day. Images were obtained using an Olympus UZ500 camera placed on a tripod over a standard photographic grey card (18% JJC neutral card) with a scale and marking plate. Four separate white fluorescent lamps were used in order to avoid shadows. Four photographs were taken of each lizard (dorsal, ventral and both lateral sides), the lizard being released without immobilisation in front of the camera and recovered between each photograph. Before each photograph, the camera was set to white balance and the resultant images were saved in TIFF format. In 2018, all lizards caught were photographed on site using a Canon EOS400D camera with Canon EF28-90mm lens on a tripod, then immediately released. In this case, the lizards were hand-held on a photographic grey card (18% JJC neutral card) with a scale and marking plate, the camera being located in shadow under vegetation in order to minimise the effect of differing lighting conditions. Before photographing, the camera was set to white balance and the resultant images were saved in RAW format. These two approaches (i.e. photography under standardised laboratory lighting and photography outside in shadow) allowed us to compare the influence of lighting on the stability of colour measurements from the photographs. For the purposes of this study, we use photographs of both lateral sides of the lizards.

### Image analysis

It is known that colour representation by reflected light depends strongly on the illumination and physical characteristics of the receivers (Wyszecki and Stiles 2000; Barnard and Funt 2002; Hill and McGraw 2006). Hence, we used a standard photographic grey card as a background in all photographs in order to minimise differences in lighting and camera setups. Further, the colours were linearly transformed in each image based on an average of 10 random pixels located on the grey card (Stevens et al. 2007; Thornbush 2008). We then cropped the standardised images to a rectangle between the front and hind legs of the lizards and transformed the RGB colour space to CIELAB, which enhances perceptive differences between colours, using D65 reference white. The images were then saved in lossless TIFF format.

We selected three pixels for each region on the cropped images where colour was to be measured. The pixels were dorsoventrally equidistant and spaced at 20, 50 and 80% of the image width in a craniocaudal direction (Fig. 1). We enlarged the area centred at each pixel to 9 × 9 pixels for 81 technical replicates of each measurement per image. The pixels for technical replicates were utilised in two alternative ways. First, we evaluated the colour of sand lizards from point measurements by calculating the average of 5 × 5 pixels centred on each technical replicate pixel for each colour channel. For the second method, we used the pixels in the dorsal, central and caudal regions as starting points for *k*-means clustering. We added a random jitter ≈ 10^−6^ to the colour channel starting points for numerical stability.

**Figure 1:**
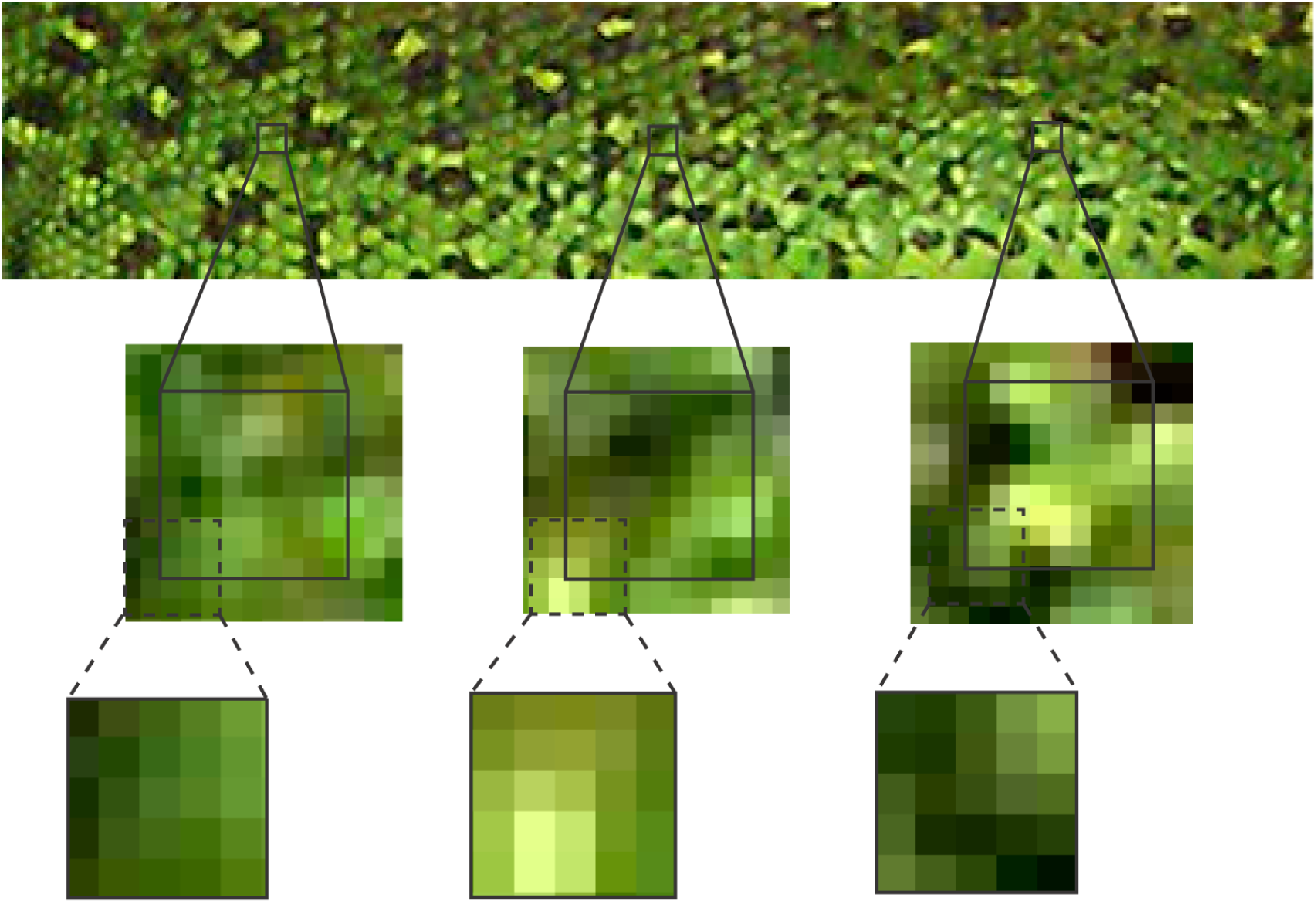
Colour assessment scheme for point measurements and starting points for *k*-means clustering. We selected three regions on the lizard’s body located equidistant laterally and at 20, 50 and 80% of the image width craniocaudally. For the point measurements, we averaged colours across the three regions, taking an area of 5 × 5 pixels in each region (dashed lines). For *k*-means clustering, we used the colours of the three pixels, one from each squared region, as starting points for the clustering algorithm. We then repeated the analysis with a sliding window approach spaced 1 pixel across a 9 × 9 area for each individual (solid lines), obtaining 81 technical replicates per photograph. These were used to assess the stability of the colour estimated using the given approach.

### Statistical analysis

For each image, we calculated the mean and standard deviation of the mean (SD) for the colour channel values. This was done for all point measurement replicates and for cluster centroids ordered by luminance. We then compared the colour means obtained for each channel under the two lighting schemes using the *t*-test, correcting the significance for multiple comparisons with the false discovery rate (FDR).

We compared the colours estimated from the point measurements to colours in the clusters using Δ*E*_00_ colour difference. The Δ*E*_00_ distance is the Euclidean distance between two colours in the CIELAB colour space, corrected for perceptive differences (Hunt 2004). All analyses were run in the R statistical platform (R Core Team 2019) using custom scripts based on the packages EBImage (Pau et al. 2010), imager (Barthelme 2019), RColorBrewer (Neuwirth 2014), scatterplot3d (Ligges and Mächler 2003), spacesXYZ (Davis 2018), stringr (Wickham 2018) and tiff (Urbanek 2013).

### Ethics statement

Sampling was based on permits 2579/2007-2.1 and 1323/527/05-5.1, issued by The Ministry of the Environment of the Slovak Republic, and JMK 38000/2018, issued by the Regional Authority of the South Moravian Region, Brno. Animal handling complied with Czech Law No. 114/1992 on Nature and Landscape Protection. The authors were authorised to handle wild lizards according to the Certificate of Professional Competence (Nos. CZ01287 and CZ03799; §15d, Act No. 246/1992 Coll.).

## Results and discussion

During the 2007 season, 167 lizards were caught, including 159 adults (79♂, 80♀) and 8 subadults (4♂, 4♀), 36 of which were re-traps. No juveniles were caught in 2007. In 2018, we caught 80 lizards, including 58 adults (21♂and 37♀), 4 subadults (2♀, 2 no sex identified) and 18 juveniles (no sex identified), 19 of which were re-traps. There was no significant difference in size between the sites, with adults measuring ♂*L* ∈ [6.0, 8.5] cm, ♀*L* ∈ [6.0, 8.7] cm and subadults ♂*L* ∈ [4.7, 4.8] cm, ♀*L* ∈ [4.9, 5.6] cm.

As a result of the lizards not being immobilised during photography in 2007, photographs of 34 lizards had to be removed from the study as body position prevented the images from being cropped to the designated rectangle.

When using pixels within a designated area as point measurements of colour, the mean colour of each lizard photograph had a greater SD than the mean colour of cluster centroids, where the pixels represented starting values in the clustering algorithm (Fig. 2A). In effect, the clustering of all pixels in the standardised image extracted the predominant colours displayed on the photographed animal, irrespective of the colour pattern. Our data showed that cluster centroids displayed lower variation in the resulting predominant colours than averaging across multiple pixels at a point measurement where the starting points shift across the lizards pattern (Fig. 2A). Point measurements of the reflectance spectra, here reduced to the red, green and blue channels, necessitates averaging across an area of 1-2mm^2^, or across multiple pixels. The average did not correspond to the colour observed on the individual when the measured area included partial colour pattern similarly as was observed in another lizards, *Zootoca vivipara* (Martin et al. 2013). To reduce such variability in the point measurements, and to increase reproducibility of the measurement, one must choose between consistency in selecting morphologically homologous areas on the body and consistency in the colours measured with respect to the colour pattern. The user must then select pixels of the intended colour, thereby bringing subjectivity into the data acquisition. Interestingly, while we found no difference in colour cluster variation between the two lighting schemes, the SD of colour channel means from point measurements in photographs taken under natural light and standardised to a grey card were significantly lower than those taken under controlled illumination in the laboratory (*t*-test: *p* < 0.05, Fig. 2B).

**Figure 2:**
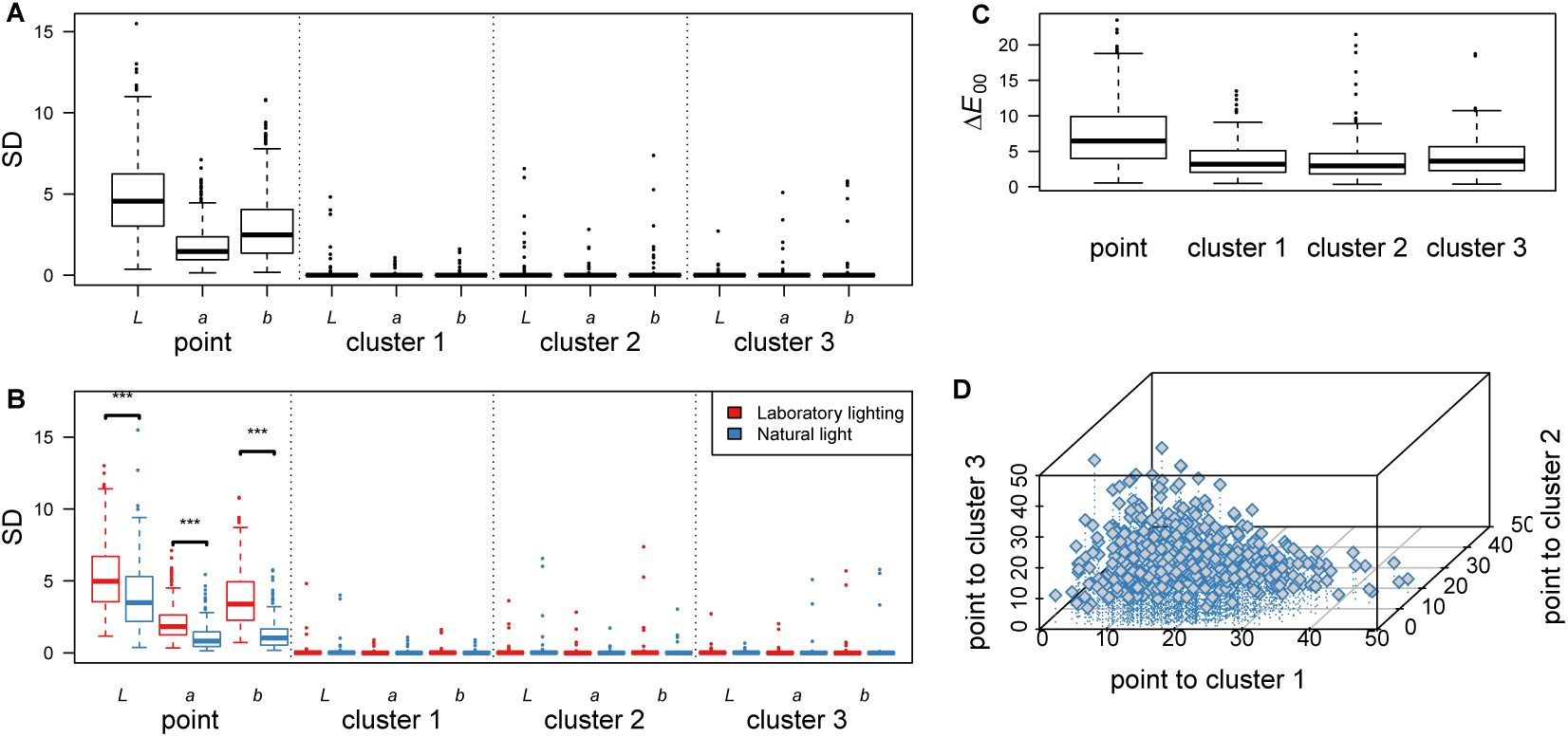
Colour measurement variation along the flanks of sand lizards. **A**. Standard deviation (SD) of colour channels in CIELAB colour space for colour means of sand lizards. Using three regions on the cropped image of a sand lizard, we averaged the colour for the point measurements and used the colours in each region as starting values for three clusters in the *k*-means clustering algorithm. **B**. SD of colour channels differentiated according to photographic lighting. The statistical significance of differences in lighting was evaluated with the *t*-test and corrected for multiple comparisons with FDR. **C**. Δ*E*_00_ colour difference between colour means estimated from the left and right flank of each animal. **D**. Δ*E*_00_ colour difference between colour means estimated from the point measurements and each of the three colour clusters. SD - standard deviation of the mean, *L -* luminance, *a* - green to red, *b -* blue to yellow, *** - *p* < 0.001.

Next, we were interested in whether colours differed on the two flanks of the lizard. Using point measurements, colour difference between paired sides of the same lizard was Δ*E*_00_ ≈ 7.4 (Fig. 2C), which would be expected in animals with colour patterns that differ laterally (Swaddle and Cuthill 1994). Colour pattern symmetry is one of the honest signals reflecting fitness of the individual, and lateral variations in colour pattern in our data influenced point measurements of colour considerably. In comparison, the colour difference between each colour cluster from the left and right side of each animal was significantly lower than that using point measurements (Δ*E*_00_ ≈ 3.9, *t*-test: *p* < 0.05). Larger differences in colour from point measurements of the two flanks of sand lizards and small differences in colours from clustering analysis of the flanks indicate that asymmetry in colour pattern is more pronounced than asymmetry in individual colours displayed on each flank of an animal.

One might ask what a point measurement of colour actually estimates in animals with a colour pattern. Intuitively, one would expect that either a colour from a point measurement represents the mean of colours contributing to the colour pattern or that it reflects the most frequent colour present on the animal. In our data, the mean colour difference between point measurements and cluster 2 was lowest with Δ*E*_00_ ≈ 4.9 (Fig. 2D) while the mean difference between clusters 1 and 3 was greater than 20. Cluster 2, as ordered by luminance of the cluster centroids, was the cluster to which most pixels were usually allocated (mean: 41.28%, range ∈ [23.3, 65.0]%). The observed relationship, i.e. where the difference between the point measurement and cluster centroid decreased as the cluster size increased, holds for all clusters, with in linear models showing significant negative slopes for all comparisons. Averaging across multiple point measurements reflected the most frequent colour.

Measuring colour in animals with complex colour patterns is challenging, with the errors associated with point measurement increasing with increasing complexity of the colour pattern. While point measurements might be appropriate for animals with large patches of similar colour, such as the green lizard (*Lacerta viridis*), for others we recommend a method of colour analysis that directly considers colour pattern. Ideally, one would combine depth of information from reflectance spectra with clustering of similar colours, as suggested here. Recent technological advances have made systems such as hyperspectral cameras more accessible for biological applications (Tedore and Nilsson 2019). Under such a system, the target object is scanned with a hyperspectral camera that provides each pixel in the image with a full reflectance spectrum. Subsequent analysis of the data cube uses image segmentation methods (i.e Coleman and Andrews 1979; Maia et al. 2019) to provide detailed information on the animal’s colour pattern, respecting its full splendour.

## Acknowledgements

We thank Adam Konečný for allowing us access to his sampling site, and Zuzana Dolinay, Markéta Harazim, Pavol Hiadlovský and Alexandra Zahradníková jr. for help with field work. Access to computing and storage facilities owned by parties and projects contributing to the National Grid Infrastructure MetaCentrum, provided under the programme “Projects of Large Research, Development, and Innovation Infrastructures” (CESNET LM2015042), is greatly appreciated. This study was supported by the Institute of Vertebrate Biology of the Czech Academy of Sciences (Grant No. 900623).

## Author contributions

TD, RS, ZH and NM conceptualised the study; TD, RS, ZH, MD and NM collected material; TD, RS, MD and NM analysed the data; TD, RS and NM wrote the manuscript, which all authors reviewed.

## References

C. Andres, F. Franke, C. Bleidorn, D. Bernhard, and M. Schlegel. Phylogenetic analysis of the *Lacerta agilis* subspecies complex. Systematics and Biodiversity, 12:43–54, 2014. doi: 10.1080/14772000.2013.878000.

K. Barnard and B. Funt. Camera characterization for color research. Color Research & Application, 27:152–163, 2002. doi: 10.1002/col.10050.

S. Barthelme. imager: Image Processing Library Based on ‘CImg’, 2019. URL https://CRAN.R-project.org/package=imager. R package version 0.41.2.

W. Bischoff. *Lacerta agilis* Linnaeus 1758 - zauneidechse. In W. Bohme, editor, Handbuch der Reptilien und Amphibien Europas, Band 2/1 Echsen II (Lacerta). Aula Verlag, 1984.

W. Bischoff. Zur Verbreitung und Systematik der Zauneidechse, *Lacerta agilis* Linnaeus, 1758. Mertensiella, 1:11–30, 1988.

G. B. Coleman and H. C. Andrews. Image segmentation by clustering. Proceedings of the IEEE, 67(5):773–785, 1979. doi: 10.1109/PROC.1979.11327.

G. Davis. spacesXYZ: CIE XYZ and some of Its Derived Color Spaces, 2018. URL https://CRAN.R-project.org/package=spacesXYZ. R package version 1.0-4.

E. Font and G. Pérez i de Lanuza. Ultraviolet reflectance of male nuptial colouration in sand lizards (*Lacerta agilis*) from the pyrenees. Amphibia-Reptilia, 28:438–443, 2007.

G. E. Hill and K. J. McGraw. Bird coloration. Harvard University Press, Cambridge, Mass., 1 edition, 2006. ISBN 0674021762.

R. Hunt. The reproduction of colour. John Wiley & Sons, Ltd, Chichester, UK, 6 edition, 2004.

S. Johnsen. How to measure color using spectrometers and calibrated photographs. Journal of Experimental Biology, 219:772–778, 2016. doi: 10.1242/jeb.124008.

T. I. Kotenko and Y. Y. Sviridenko. Variability of coloration and pattern of the sand lizard, *Lacerta agilis* (Reptilia, Sauria, Lacertidae): Methodic aspects. Vestnik zoologii, 44:137–162, 2010. in Russian.

U. Ligges and M. Mächler. Scatterplot3d - an R package for visualizing multivariate data. Journal of Statistical Software, 8(11):1–20, 2003. URL http://www.jstatsoft.org.

R. Maia, H. Gruson, J. A. Endler, and T. E. White. pavo 2: New tools for the spectral and spatial analysis of colour in R. Methods in Ecology and Evolution, 10:1097–1107, 2019. doi: 10.1111/2041-210X.13174.

M. Martin, S. Meylan, D. Gomez, and J.-F. Le Galliard. Ultraviolet and carotenoid-based coloration in the viviparous lizard *Zootoca vivipara* (Squamata: Lacertidae) in relation to age, sex, and morphology. Biological Journal of the Linnean Society, 110:128–141, 2013. doi: 10.1111/bij.12104.

D. E. McCoy and R. O. Prum. Convergent evolution of super black plumage near bright color in 15 bird families. Journal of Experimental Biology, 222, 2019. doi: 10.1242/jeb.208140.

E. Neuwirth. RColorBrewer: ColorBrewer Palettes, 2014. URL https://CRAN.R-project.org/package=RColorBrewer. R package version 1.1-2.

G. Pau, F. Fuchs, O. Sklyar, M. Boutros, and W. Huber. EBImage: an R package for image processing with applications to cellular phenotypes. Bioinformatics, 26:979–981, 2010. doi: 10.1093/bioinformatics/btq046.

R Core Team. R: A Language and Environment for Statistical Computing. R Foundation for Statistical Computing, Vienna, Austria, 2019. URL https://www.R-project.org/.

F. Rohlf. tpsDig, Digitize Landmarks and Outlines, Version 2.05, 2005. URL http://life.bio.sunysb.edu/morph/. Accessed 21 Sept 2012.

M. Stevens, C. A. Párraga, I. C. Cuthill, J. C. Patridge, and T. S. Tros-cianko. Using digital photography to study animal coloration. Biological Journal of the Linnean Society, 90:211–237, 2007. doi: 10.1111/j.1095-8312.2007.00725.x.

J. Swaddle and I. Cuthill. Preference for symmetric males by female zebra finches. Nature, 367:165–166, 1994. doi: 10.1038/367165a0.

C. Tedore and D. Nilsson. Avian UV vision enhances leaf surface contrasts in forest environments. Nature Communications, 10:238, 2019. doi: 10.1038/s41467-018-08142-5.

G. Testylier and P. Gourmelon. Spectrophotometry in vivo, a technique for local and direct enzymatic assays: application to brain acetyl-cholinesterase. Proceedings of the National Academy of Sciences, 84:8145–8149, 1987. doi: 10.1073/pnas.84.22.8145.

M. J. Thornbush. Grayscale calibration of outdoor photographic surveys of historical stone walls in oxford, england. Color Research & Application, 33 (1):61–67, 2008. ISSN 03612317. doi: 10.1002/col.20374.

S. Urbanek. tiff: Read and write TIFF images, 2013. URL https://CRAN.R-project.org/package=tiff. R package version 0.1-5.

M. Šulc, J. Troscianko, G. Štětková, A. E. Hughes, V. Jelínek, M. Capek, and M. Honza. Mimicry cannot explain rejection type in a host–brood parasite system. Animal Behaviour, 155:111–118, 2019. doi: https://doi.org/10.1016/j.anbehav.2019.05.021.

A. Waichman. An alphanumeric code for toe clipping amphibians and reptiles. Herpetological Review, 23:19–21, 1992.

H. Wickham. stringr: Simple, Consistent Wrappers for Common String Operations, 2018. URL https://CRAN.R-project.org/package=stringr.

G. Wyszecki and W. S. Stiles. Color science: concepts and methods, quantitative data, and formulae. John Wiley & Sons, New York, 2000.

